# Intragenic Complementation of *Cellulose Synthase-Like D1* alleles in Root Hair Development

**DOI:** 10.1101/2020.03.02.973545

**Authors:** Bogumil J. Karas, Loretta Ross, Mara Novero, Lisa Amyot, Sayaka Inada, Michiharu Nakano, Tatsuya Sakai, Sushei Sato, Jeremy Dale Murray, Paola Bonfante, Krzysztof Szczyglowski

## Abstract

Root hair cells form the primary interface of plants with the soil environment, playing key roles in nutrient uptake and plant defense. In addition, they are typically the first cells infected by nitrogen-fixing soil bacteria during the root nodule symbiosis. Here we report a role for the *Cellulose Synthase-Like D1* (*CSLD1*) gene in root hair development in *Lotus japonicus*. CSLD1 belongs to the cellulose synthase protein family that includes cellulose synthases, and cellulose synthase-like proteins, the latter thought to be involved in the biosynthesis of hemicellulose. We describe 11 *csld1* mutant alleles that have either short (*Ljcsld1-1*) or variable length roots hairs (*Ljcsld1-2* to *11*). Examination of *Ljcsld1-1* and one variable-length root hair mutant, *Ljcsld1-6* showed increased root hair cell wall thickness, which in *Ljcsld1-1* was more pronounced, suggesting a possible link with the defect in root nodule symbiosis. In addition, *Ljcsld1-1* heterozygotes had intermediate root hair lengths, between those of wild type and the homozygotes. Intragenic complementation was observed between alleles with mutations in the N-terminal domain and other alleles, suggesting modularity of CSLD1 function and that it may operate as a homodimer or multimer.

**One sentence summary:** This research describes novel gain- and loss-of-function mutations at the *Lotus japonicus CELLULOSE SYNTHASE-LIKE D1* locus and analyzes their impact on root hair development.

## Introduction

Plant cell walls constitute a dynamic yet rigid interface between cells or between the cell and the external environment. While providing structural support and protection, they also act as filtering structures. Composed primarily of cellulose, hemicellulose, and pectin (Lampugnani et al., 2018), cell walls vary considerably based on plant species as well as tissue type (Popper et al., 2014; Höfte and Voxeur, 2017; Penning et al., 2019). How each cell wall component is synthesized is still not fully understood.

Most of the research aimed at identifying the various players required for cell wall synthesis has been performed in the model plant *Arabidopsis thaliana* (Liepman et al., 2010). Cellulose, the main structural component of plant cell walls, is synthesized by large, plasma membrane-localized complexes consisting of cellulose synthase (CESA) subunits. Genome sequencing has revealed that in *A. thaliana* there are ten genes that encode CESAs (Richmond, 2000). While CESA1, CESA3 and a CESA6-like protein (CESA2, CESA5, CESA6 or CESA9) are required for primary cell wall synthesis (Arioli et al., 1998; Fagard et al., 2000; Scheible and Pauly, 2004; Desprez et al., 2007; Persson et al., 2007; Lampugnani et al., 2019), a complex containing CESA4, CESA7, and CESA8 subunits is required for secondary cell wall formation (Taylor et al., 1999; Taylor et al., 2000; Taylor et al., 2003). These functional roles were supported by a global expression study which showed that *CESA1, CESA3*, and *CESA6* are all highly and ubiquitously expressed, whereas *CESA4, CESA7*, and *CESA8* are mainly expressed in stems suggesting they may be required for the formation of the secondary cell wall (Hamann et al., 2004).

A family of related cellulose synthase-like (CSL) proteins was identified based on a conserved β-glycosyltransferase D, D, D, QXXRW motif (Saxena et al., 1995; Richmond and Somerville, 2000). Arabidopsis has 30 *CSL* genes which have been classified into six groups: *CSLA, B, C, D, E*, and *G*. Two additional *CSL* groups, *CSLF* and *CSLH*, have been identified in grasses (Lerouxel et al., 2006) as well as a third group, *CSLJ*, which is present in grasses and in some dicots (Yin et al., 2009). Compared to CESAs, the function of CSLs is less well studied and based on their diverse expression patterns, they are thought to play more specialized roles. Several studies have indicated that CSLs are involved in the synthesis of hemicellulose, which is a heterogeneous group of polysaccharides that provide additional structural support to the cell wall through their interactions with cellulose. For instance, *Arabidopsis* CSLAs have been shown to have mannan and glucomannan synthase activity (Liepman et al., 2005; Goubet et al., 2009), while it has been demonstrated that CSLCs are involved in the synthesis of the xyloglucan backbone (Cocuron et al., 2007). Verhertbruggen et al. (2011) also demonstrated that CSLD3 has mannan synthase activity, along with co-expressed CSLD2 and CSLD5 (Verhertbruggen et al., 2011), while Park and colleagues (Park et al., 2011) showed that chimeric CSLD3 containing the CESA6 catalytic domain could complement the *csld3* mutant.

Of the CSLs, the CSLD group shares the highest similarity with CESAs (Richmond and Sommerville, 2011). Studies on *csld* mutants have suggested a role for these genes in stem and tip cell growth, as well as cellulose deposition. Thus far, mutations in *CSLD5* have been associated with defects in stem growth (Bernal et al., 2007), in *CSLD1* and *CSLD4* with pollen tube development (Bernal et al., 2008; Wang et al., 2011), and in *CSLD2* and *CSLD3* with defective root hairs (Favery et al., 2001; Wang et al., 2001; Bernal et al., 2008; Galway et al., 2011). Yoo et al. (Yoo et al., 2012) also showed that plants lacking both CSLD2 and CSLD3 had defects in female gametophyte development.

Root hair cells are large, rapidly growing, and easily observable, making them an ideal system for studying cell wall development (Foreman and Dolan, 2001). We have identified several root hair mutants from a screen for genetic suppressors of the *Lotus japonicus har1-1* hypernodulation phenotype (Murray et al., 2006b; Murray et al., 2006a), where developmental defects in the root epidermis led to the impairment of root nodule symbiosis. Our initial microscopic observations, as well as genetic crosses, allowed us to classify these root hair mutants into four complementation groups: *hairless*, corresponding to the *L. japonicus Root Hairless* locus, (*LjRHL*), *petite* (*L. japonicus Petite Root Hairs, LjPRH*), *short* (*L. japonicus Short Root Hairs, LjSRH*) or *variable* (*L. japonicus Variable Root Hairs, LjVRH*) (Karas et al., 2005). The *LjRHL* locus was identified as a basic helix-loop-helix protein, and its orthologues *AtLRL1, AtLRL2*, and *AtLRL3* were shown to be redundantly required for root hair development in *Arabidopsis* (Karas et al., 2009).

In this study, we focused on *L. japonicus srh* and *vrh* developmental mutants. Map-based cloning revealed that both short and variable root hair mutant lines carried lesions in the *CSLD1* gene, encoding a protein with the highest homology to members of the *Arabidopsis* CSLD family. The root hairs of *csld1* mutants had thicker cell walls, which in *Ljcsld1-1* was particularly pronounced and was associated with impaired colonization of roots by *M. loti*. Unexpectedly, a subset of our inter-allelic crosses resulted in the restoration of wild type root hairs (intragenic complementation). Thus, our allelic series provides a useful model for studying how CSLD proteins interact with each other but, and how this influences the structure of the root hair cell wall.

## Results

### Identification of 11 *LjCSLD1* alleles

The identification and phenotypic characterization of three allelic *L. japonicus* mutant lines of the variable root hair phenotype (*Ljvrh1-1, Ljvrh1-2*, and *Ljvrh1-3*) and one additional line that showed a short root hair phenotype (*Ljsrh1*) were described earlier (Karas et al., 2005). By surveying our in-house collection of *L. japonicus* mutants and the NBRP Legume Base resource (https://www.legumebase.brc.miyazaki-u.ac.jp/), an additional 26 lines with altered root hair phenotypes were identified. While none of these lines had short root hairs, 15 showed phenotypes resembling *Ljvrh1* (Fig. 1A).

**Figure 1.**
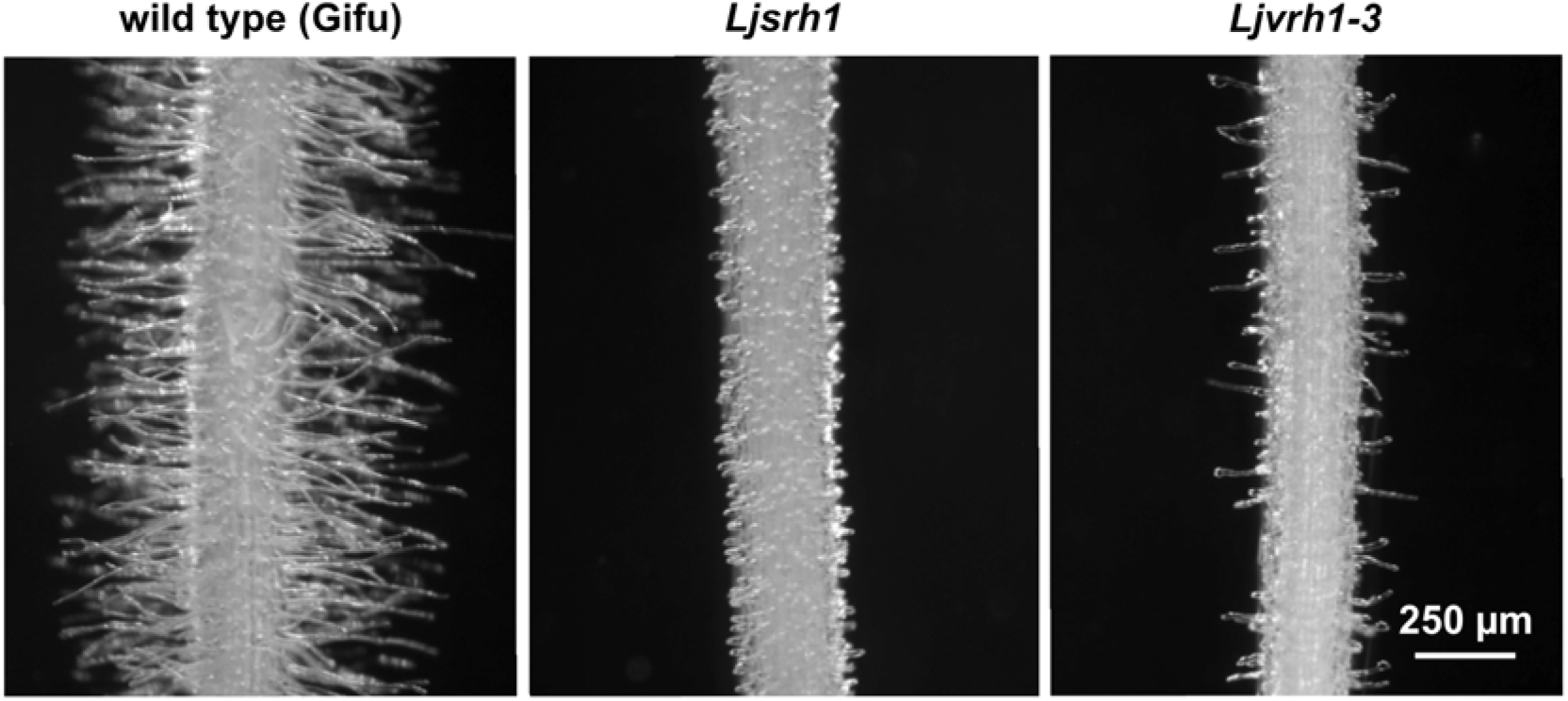
Root hair phenotypes. Root hairs of the wild type *L. japonicus* Gifu and *Ljcsld1-1* (initially identified as *Ljsrh1*) and *Ljcsld1-2* (initially identified as *Ljvrh1-1*) mutant lines are shown. Root segments were photographed at approximately 0.5-1 cm from the root tip, where fully elongated root hairs are present in Gifu (see Karas et al., 2005). Note differences in root hair length between the three lines. The 250 µm scale applies to all three images.

A map-based cloning approach was undertaken in order to identify the underlying genetic lesions. The 0.8 cM genetic interval, which was previously defined to contain the *LjVRH1* and *LjSRH1*, as delineated by the TM1419 and TM0127 flanking markers (Karas et al., 2005), encompassed five overlapping TAC clones (Supplemental Fig. S1). A survey of genes present on these clones identified, among others, a *Cellulose Synthase-Like D* gene (*LjCSLD1*) that was located on TM0757 (Supplemental Fig. S1). This gene was considered as a viable candidate for either *LjVRH1* or *LjSRH1* locus, as proteins belonging to the *Arabidopsis* AtCSLD subfamily, such as KOJAK, were shown to be required for root hair cell morphogenesis (Favery et al., 2001). Sequencing of the predicted coding region of *LjCSLD1* in wild type and the three *Ljvrh* mutants identified a C3494T transition in *Ljvrh1-1*, while *Ljvrh1-2* and *Ljvrh1-3* carried C3939T and G530A transitions, respectively. The same locus was amplified and sequenced from the 15 additional *L. japonicus* mutant lines with the *Ljvrh-like* phenotypes. Seven of these carried single nucleotide substitutions in the predicted coding region of the *LjCSLD1* gene (Table S1) while the remaining lines had the wild-type *LjCSLD1* sequence, suggesting that their mutant root hair phenotypes were determined by mutations in an independent locus or loci.

A parallel effort to map-based clone the *LjSRH1* locus was also undertaken. The initial genetic complementation analysis yielded F1 plants with wild-type root hairs, suggesting that *LjSRH1* was independent from *LjVRH1*. However, mapping experiments localized both loci to the same 0.8 cM genetic interval at the bottom of *L. japonicus* chromosome III (Karas et al., 2005). The position of the *Ljsrh1* mutation was further delimited to the 50 kb region between flanking markers JM010 and JM003 (Supplemental Fig. S1). This region contained eight predicted genes, including *LjCSLD1* (Supplemental Fig. S1). Sequencing of all eight genes from wild-type and the *Ljsrh1* mutant identified a single nucleotide substitution, C220T, in *LjCSLD1*, while the nucleotide sequence of the remaining seven genes was wild type. The *Ljsrh1* allele was tentatively renamed as *Ljcsld1-1*. The same *Ljcsld* nomenclature was used for the variable root hair phenotype-associated mutations, with allele numbering reflective of the relative position of a given mutation along the *LjCSLD1* gene sequence (Table S1).

### *Lotus LjCSLD1* and *Arabidopsis AtCSLD2* and *AtCSLD3* are functionally conserved

To confirm that the mutations at the *LjCSLD1* locus were causative to both short and variable root hair phenotypes, *in planta* complementation experiments were performed. A binary vector containing an 8.5 kb genomic fragment encompassing the entire *LjCSLD1* locus was introduced into roots of *Ljcsld1-1, Ljcsld1-2* and *Ljcsld1-6* mutant lines by *Agrobacterium rhizogenes*-mediated transformation (Murray et al., 2007). The resulting transgenic hairy roots produced wild type root hairs. In contrast, a control transformation, using an *A. rhizogenes* strain carrying an empty vector, failed to complement the defective root hair phenotypes (Fig. 2A).

**Figure 2.**
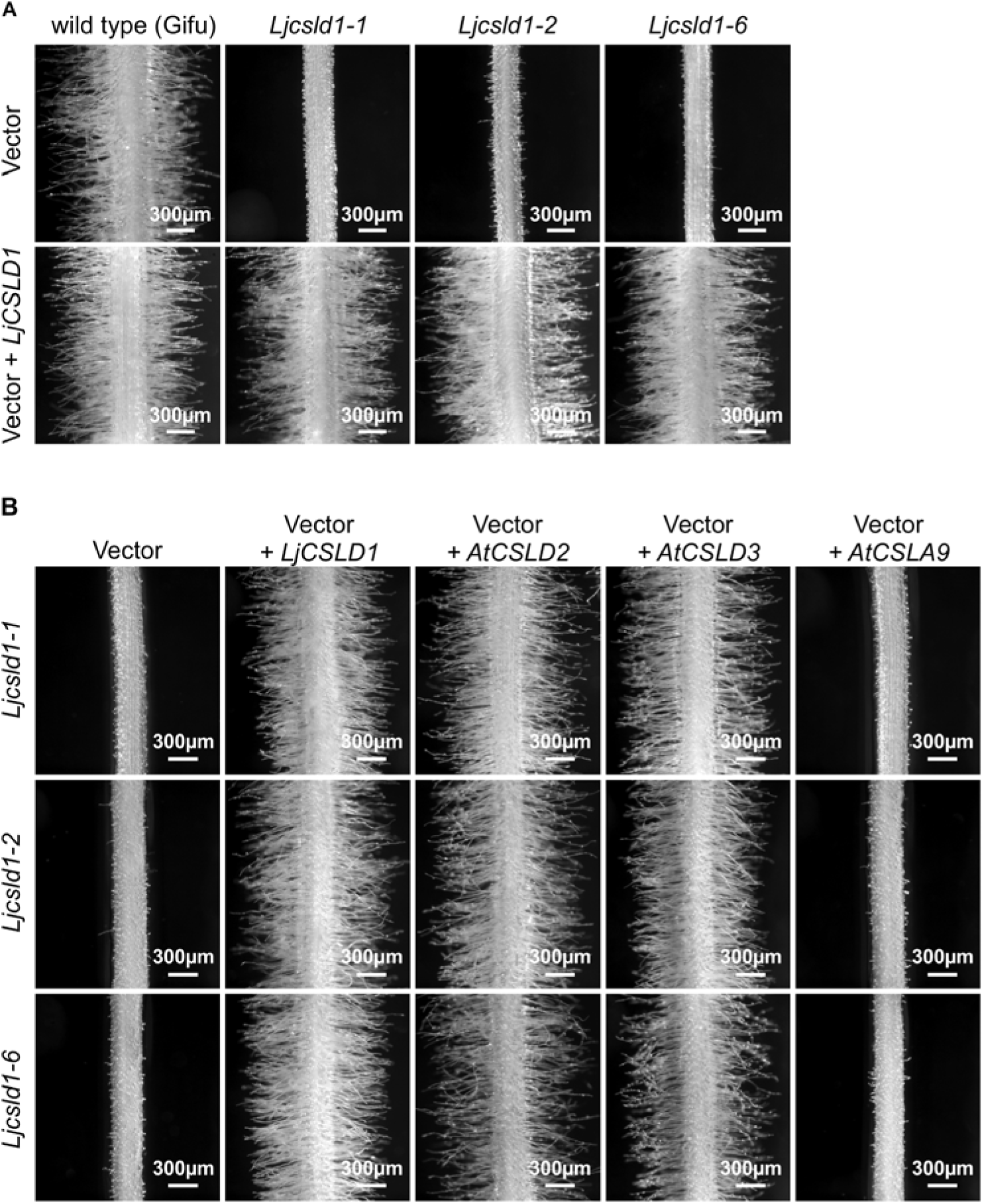
Genetic complementation. (A) The *LjCSLD1* gene complements the *Ljcsld1-1* short and *Ljcsld1-2* and *Ljcsld1-6* variable root hair phenotypes. The mutant plants were transformed by either the *A. rhizogenes* strain AR10 carrying control binary vector with no insert (vector, upper row), or the same vector with the entire genomic version of the *LjCSLD1* gene (vector + *LjCSLD1;* bottom row). (B) Arabidopsis *AtCSLD2* and *AtCSLD3* complement the short root hair phenotypes of *Ljcsld1-1* (top row) and the variable root hair phenotypes of *Ljcsld1-2* (middle row) and *Ljcsld1-6* (bottom row) while *AtCSLA9* does not. The corresponding *Arabidopsis* cDNAs were expressed in transgenic hairy roots under the control of *CaMV* 35S promoter. The empty pEarley101 binary vector, containing the 35S promoter, was used as negative control while *LjCSLD1* cDNA served as the positive control.

Given the amino acid similarity between LjCSLD1 and *Arabidopsis* AtCSLD2 and AtCSLD3 (Supplemental Fig. S2), an inter-species complementation test was performed. Expression of *AtCSLD2* and *AtCSLD3* under the control of the *CaMV* 35S promoter in transgenic hairy roots rescued the short and variable root hair phenotypes of *L. japonicus Ljcsld1-1, Ljcsld1-2*, and *Ljcsld1-6* mutants. In contrast, the more distantly related *AtCSLA9* did not complement the root hair defects of any of the three allelic lines tested (Fig. 2B). Based on these data, we concluded that the *LjCSLD1* gene, identified through map-based cloning approaches, indeed corresponded to the *L. japonicus LjVRH1/LjSRH1* locus and is functionally conserved with *Arabidopsis AtCSLD2* and *AtCSLD3*.

### Structure of the LjCSLD1 protein

We used RACE to characterise the *LjCSLD1* transcripts. Our results show that *LjCSLD1* is comprised of four exons and three introns and produces an mRNA of 3805 nucleotide (nt) in length, with a 3450 nt long open reading frame (ORF), encoding the predicted LjCSLD1 protein of approximately 129 kD. The ORF is flanked by 152 and 203 nt long 5’ and 3’ untranslated regions (UTR).

The LjCSLD1 protein contains two cysteine (C)-rich motifs of CX_4_CX_15_CXC and CX_2_CX_11_CX_2_C, starting at the amino acid residues 131 and 159, respectively, which are mostly conserved with AtCESA1 (Fig. 3 and Supplemental Fig S3A). Furthermore, four sub-domains (U1-U4), encompassing three highly conserved aspartic acid residues (D) and the QXXRW motif that are characteristic of the processive β-glucosyltransferases in plants and bacteria, were present in the presumed globular region of LjCSLD1 (Supplemental Fig. S3A). This region is flanked by eight predicted trans-membrane segments; two of these segments are present in the N-terminal portion of the region, while the remaining six transmembrane domains are in the C-terminal region (Fig. 3, Supplemental Fig. S3A).

**Figure 3.**
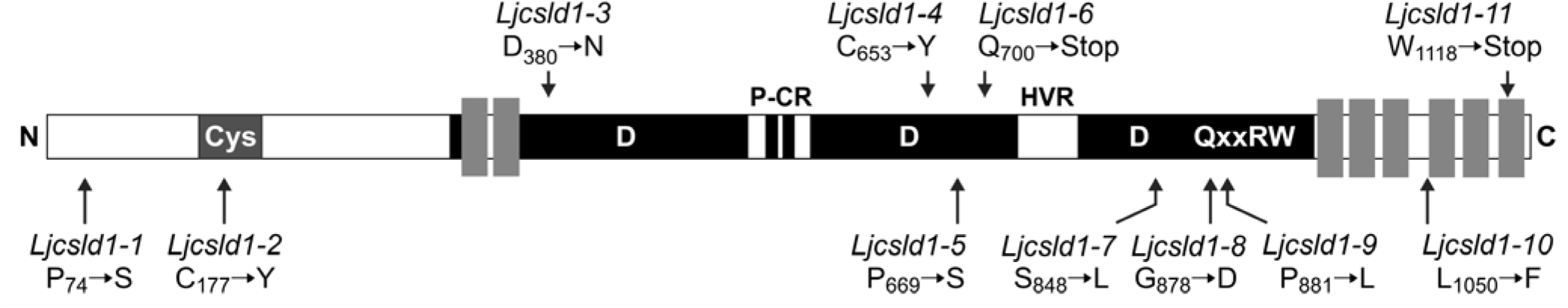
Schematic of the *LjCSLD1* protein structure showing positions of the molecular lesions for all mutant alleles. Black boxes represent conserved sequences among members of the CSLD protein family and grey rectangles represent transmembrane domains; (Cys), cysteine-rich region; (P-CR), the plant-conserved region; (HVR), plant hyper-variable region; D, D, D, QxxRW represents a conserved β-glycosyltransferase motif where D is aspartic acid; Q, glutamine; x, variable residue; R, arginine and W, tryptophan.

Two mutations, *Ljcsld1-1* and *Ljcsld1-2*, affected the LjCSLD1 N-terminal region, where substitutions of P_74_ to S and C_177_ to Y have occurred, respectively (Fig. 3, Table 1). P_74_ resides within a short proline-rich domain (PPTPD) located close to the N-terminus of LjCSLD1. This domain is highly conserved in CSLD proteins but it is absent from cellulose synthase catalytic subunit proteins (CESA) and other members of the CSL protein family. The C_177_ residue constitutes a part of the LjCSLD1 C-rich region. This region is present in all other CSLDs and has been postulated to be involved in mediating protein-protein interactions (Delmer, 1999). Seven additional mutations, *Ljcsld1-3* to *Ljcsld1-9*, were positioned within the presumed globular region of the LjCSLD1 protein (Table 1). With the exception of *Ljcsld1-6*, where a single nucleotide change of C_3494_ to T resulted in a premature stop codon, all remaining mutations were non-synonymous, leading to amino acid substitutions. Given the overall high level of amino acid sequence conservation, it was not surprising to find that all of these mutations altered amino acid residues that are conserved in CSLD proteins. Notably, however, two of these mutations, *Ljcsld1-7* and *Ljcsld1-9*, were localized within the highly conserved U3 and U4 domains of LjCSLD1. These domains are thought to participate in substrate binding and/or in mediating the catalytic activity of the protein. The two remaining mutations, *Ljcsld1-10* and *Ljcsld1-11* affected the C-terminal portion of the protein, encompassing the six transmembrane domains (Fig. 3, Table 1).

### Intra-allelic complementation at the *LjCSLD1* locus

Our initial, expectation that two independent loci were involved in the observed variable and short root hair phenotypes was based on the observation of the wild-type root hair phenotype in the F1 progeny derived from the cross between homozygous *Ljcsld1-1* (short root hair phenotype) and *Ljcsld1-2* (variable root hair phenotype). As mutations at only one locus, *LjCSLD1* was found to be causative for both phenotypes, this suggested that intra-allelic complementation must have accounted for the observed effect. We tested this assumption further by using all 11 *Ljcsld1* alleles (Fig. 4).

**Figure 4.**
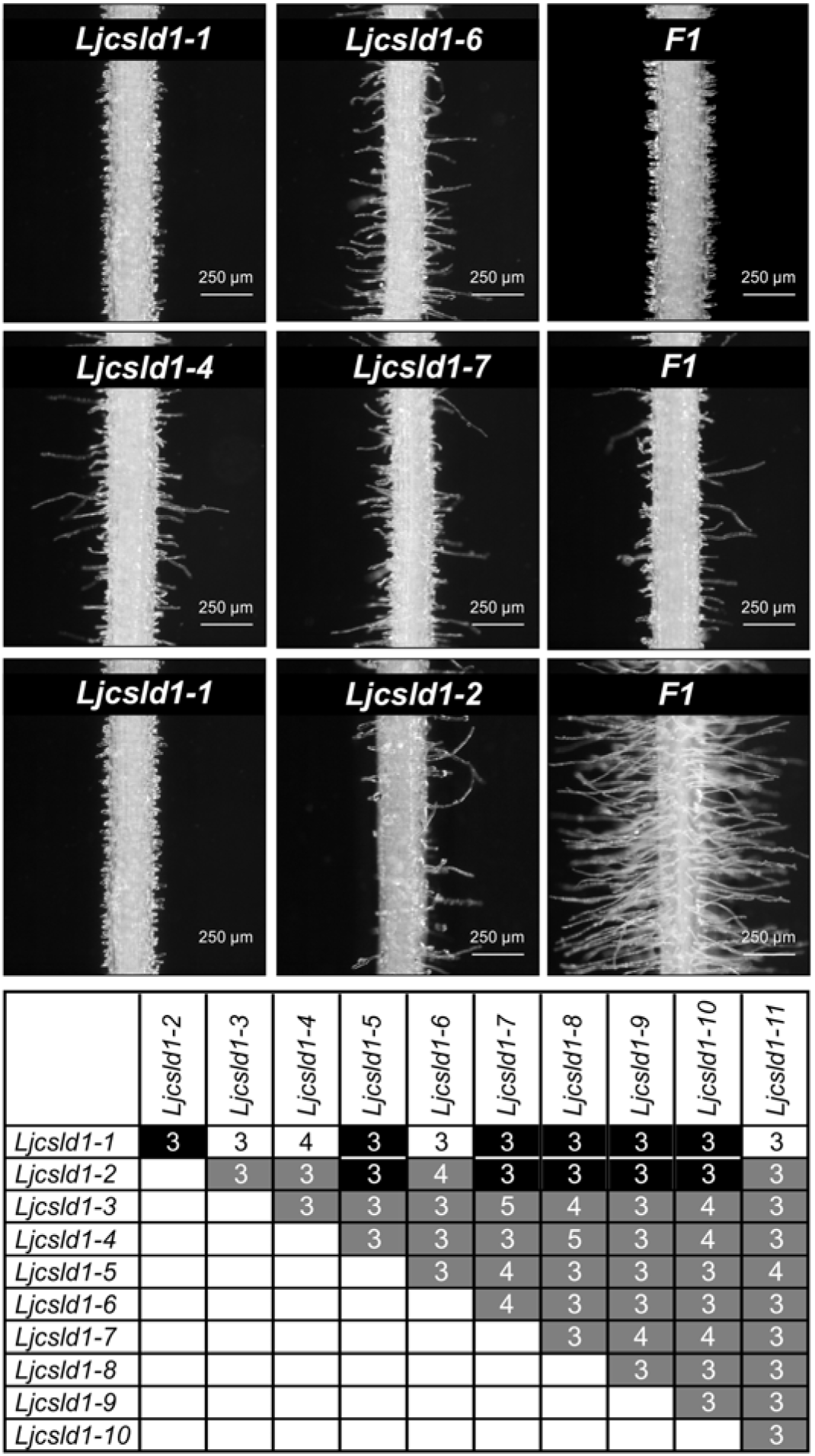
Intragenic complementation at the *LjCSLD1* locus. Allelic crosses were performed among all 11 *Ljcsld1* variants. Representative examples of three different root hair phenotypes, short (top row), variable (middle row) and wild-type (bottom row), as recovered in F1, are shown along with phenotypes of parental lines. Root segments were photographed at approximately 0.5-1 cm from the root tip. The table summarizes results of all allelic crosses; shaded boxes represent one of the three F1 root hair phenotypes: black – wild type, white – short, grey – variable root hairs. The numbers of independent crosses for each allelic combination are indicated in the shaded boxes.

As expected for crosses between different alleles of a single gene, the majority of crosses resulted in F1 progeny with mutant, short or variable root hair phenotypes (Fig. 4, see also Supplementary file 2). Interestingly, however, when either *Ljcsld1-1* or *Ljcsld1-2* were used as a crossing partner with each other or with *Ljclsd1-5* and *Ljclsd1-7* through to *Ljcsld1-10*, a wild-type like root hair phenotype was restored (Fig. 4, see also Supplementary file 2).

### Expression and functional characterization of LjCSLD1

The *LjCSLD1* mRNA was found to be present in all *L. japonicus* tissues tested, including uninoculated roots, and nodules that formed upon inoculation with *Mesorhizobium loti* (Fig. 5A). Histochemical analysis of the *LjCSLD1* gene expression in transgenic hairy roots carrying the corresponding promoter sequence fused to the coding region of the GUS reporter gene showed the localization of the promoter activity mostly in emerging to fully elongated root hairs of wild-type plants (Fig. 5B-E). These results were, therefore, consistent with the predicted role of the *LjCSLD1* gene in mediating root hair elongation in *L. japonicu*s.

**Figure 5.**
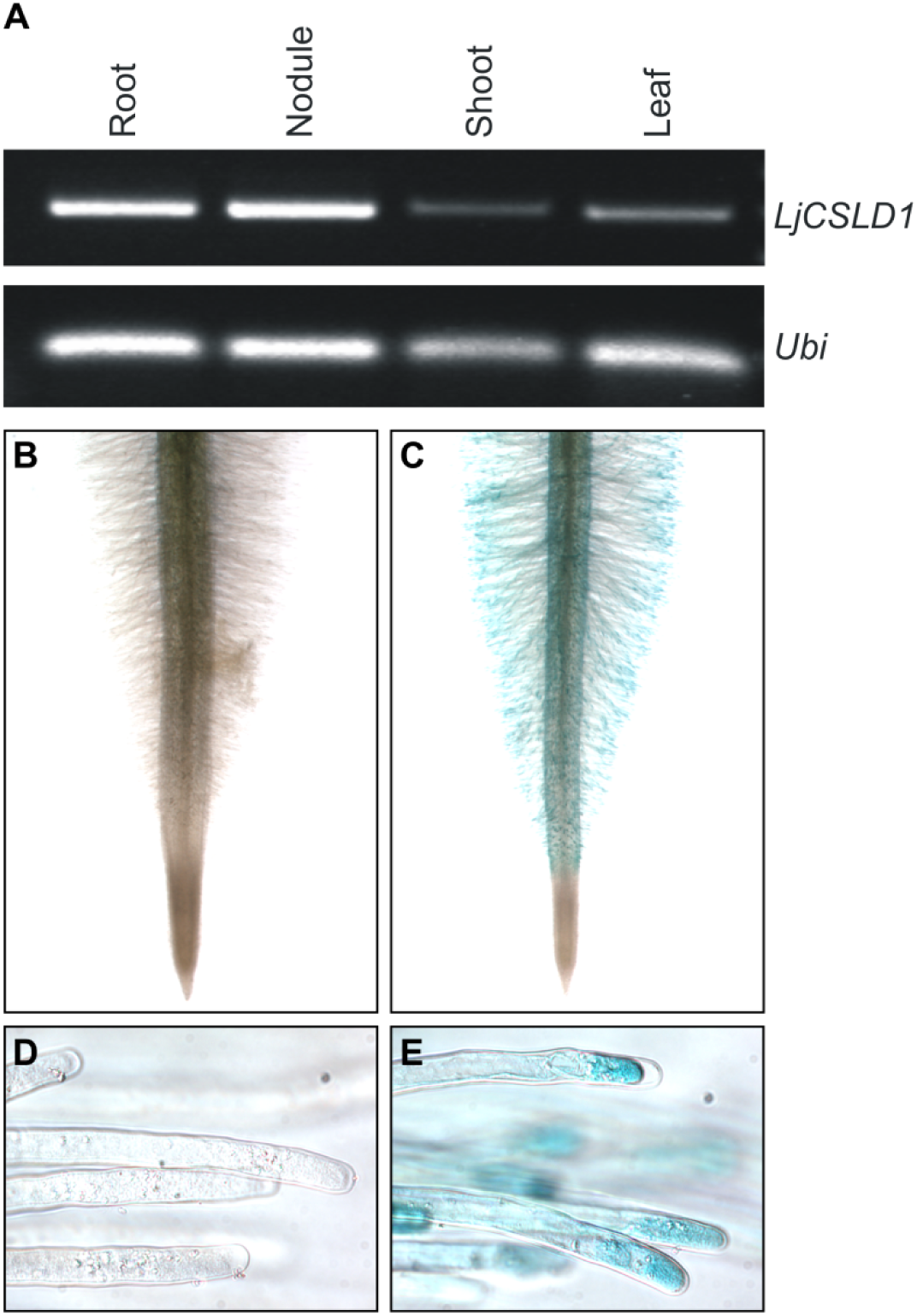
Expression of *LjCSLD1*. (A) *LjCSLD1* mRNA is present in various *L. japonicus* tissues as assayed by RT-PCR. mRNA for *Ubiquitin* (*Ubi*) was amplified and used as the RNA loading control. (B-E) The *GUS*-reporter gene activity driven by the *LjCSLD1* promoter localizes to root hair tips. Wild-type (Gifu) plants were transformed with A. *rhizogenes* strain AR10 carrying the control binary vector containing the *LjCSLD1* promoter only (B and D), or the same vector with the *LjCSLD1* promoter transcriptionally fused to the *uidA* gene coding region (C and E). The resulting hairy roots were stained for the GUS activity and photographed.

To gain insight into the biochemical function of LjCSLD1, two parallel approaches were undertaken. First, comparative chemical analyses of the root cell wall composition were performed. Although the use of only root hairs would have been preferable, this approach was not viable due to their overall scarcity in the mutant roots. Therefore, the entire roots of young wild-type and mutant seedlings were harvested and analyzed for monosaccharide content in cellulose and cell wall matrix polysaccharide fractions of the wall. In comparison with wild-type roots, glucose content in the cellulose fraction of the wall was slightly, yet significantly diminished in four mutant lines (*Ljcsld1-2*, -*4*, -*6*, and -*7*) of the variable root hair phenotype, while it was increased in *Ljcsld1-1*, the mutant line with the short root hair phenotype (Table S2A). In all mutants tested, the level of mannose was significantly decreased in the hemicellulose fraction, relative to the wild-type control. Consistent with this result, the levels of mannose, as well as galactose, were significantly lower in all mutants tested when neutral monosaccharides were quantified from an independent preparation of cell wall matrix polysaccharides (Table S2B). While the content of other neutral sugars remained unchanged in *Ljcsld1-1* in comparison to wild-type samples, the levels of fructose, arabinose, and xylose were altered in the mutants of the variable root hair phenotype. We also detected ribose which most likely results from contaminating RNA (Fry, 1988).

### Evaluation of root hair cell wall thickness

Evaluation of root hair cell wall thickness in the mature zone of the short root hair mutant (*Ljcsld1-1*), one of the variable root hair mutants (*Ljcsld1-6*), and wild type revealed that the mutants had significantly thicker cell walls than wild type. Observations performed on at least five root hairs belonging to at least five different individuals consistently demonstrated that *Ljcsld1-1* showed the largest increase (Fig. 6, Supplemental Fig. S4), with a 0.37 μm-thick cell wall when compared to the cell wall from wild-type (0.15 μm-thick). There was no difference in cell wall thickness between the short and long root hairs of *Ljcsld1-6* (Supplemental Fig. S4). When observed at higher magnification, the cell wall structure displayed dramatic differences. While the wild-type cell wall revealed the expected layered wall, where electron transparent fibrils mostly run parallel or loosely crossed (see arrows) within a loose electron-dense matrix (Emons and Ketelaar, 2009), the *Ljcsld1-1* genotype showed short and highly crossed fibrils embedded inside a dense matrix. *Ljcsld1-6* cell walls revealed an intermediate organization: in the short-type root hairs, the structure resembled that of *Ljcsld1-1* with short and crossed fibrils, while cell walls from long-type root hairs were characterized by a loose structure with bundles of long ordered fibrils. We then compared root hairs from young wild-type and *Ljcsld1-1* roots and found that the cell wall of *Ljcsld1-1* was almost three-times thicker (Fig. 6 I-L, Supplemental Fig. S5).

**Figure 6.**
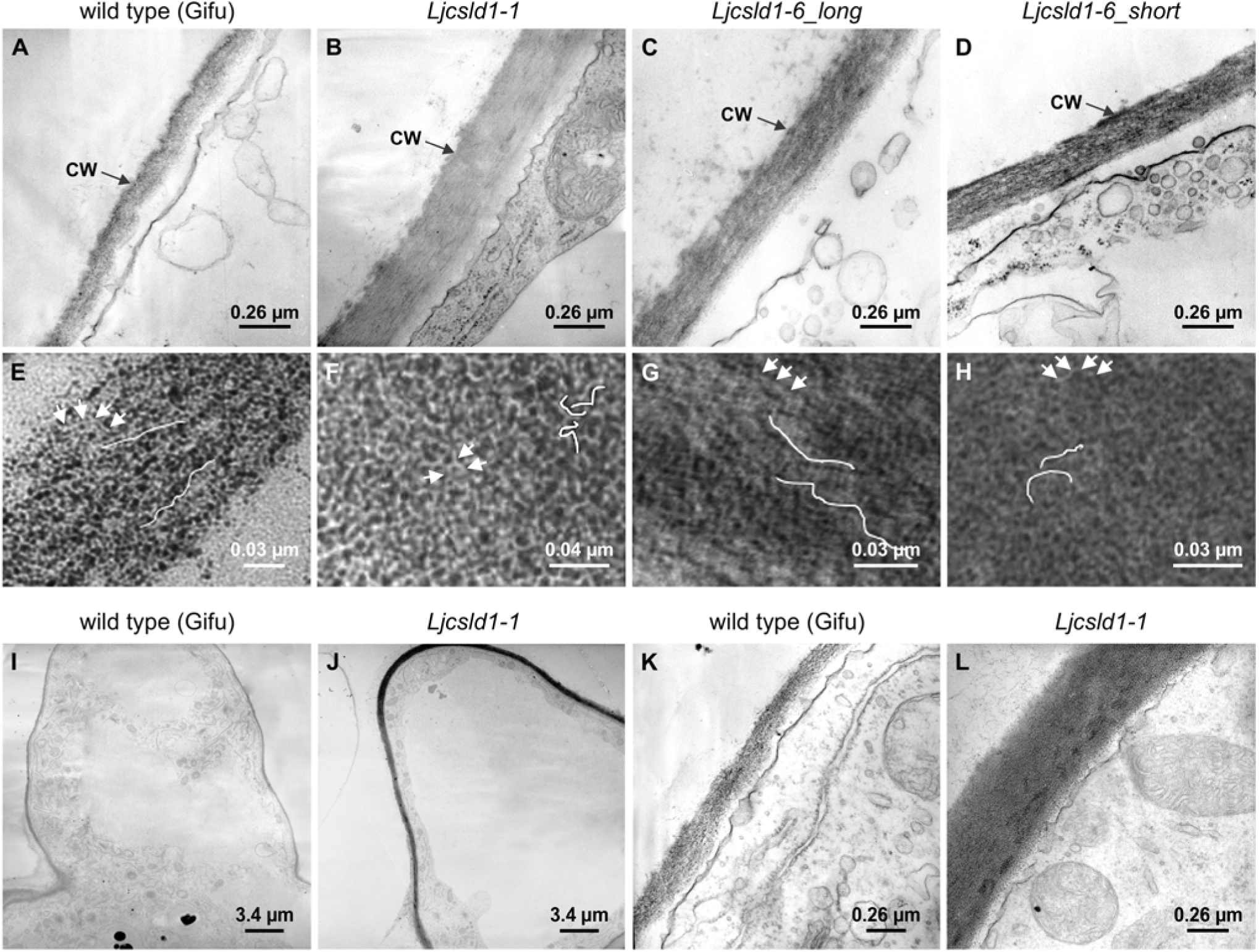
*Ljcsld1-1* and *Ljcsld1-6* alter root hair cell wall thickness and structure. (A-D) Transmission electron microscopy images of cell walls (CW) of root hairs of the mature root zone in *L. japonicus* Gifu (wild type Gifu) and *Ljcsld1-1* and *Ljsld1-6* mutants. Note that *Ljcsld1-6* displays the variable root hair phenotype; therefore, longitudinal sections for two representative types of root hairs, short and elongated (long), that are formed by *Ljcsld1-6* were analyzed. (E-H) Organization of microfibrils in cell walls is shown; wiggly lines and arrows reflect approximate shape and length of microfibrils for a given genotype and associated root hair types. (I-L) The cell wall of young *Ljcsld1-1* root hairs is almost three times thicker in comparison with the wild-type equivalent. Root hairs positioned close to the root apex, within the root hair elongation zone, were analyzed. Representative TEM images of longitudinal sections and the corresponding close-ups to root hairs cell walls in *L. japonicus* Gifu (I and K) and *Ljcsld1-1* (J and L) are shown.

### *Ljcsld1-1* has semi-dominant effect on root hair phenotype and affects root nodule development

Since the *Ljcsld1-1* allele developed thicker root hair cell walls, we decided to evaluate the phenotype of heterozygote plants in greater detail to see whether this allele exerts a semi-dominant effect on root hair growth. F2 progeny derived from the crosses between wild type and either *Ljcsld1-1* or *Ljcsld1-2* were generated and then the collective root hair length of the mature root zone (∼ 1 cm above the root tip) was measured and each plant showing wild-type-like phenotype was genotyped. There was no difference in root hair length between wild-type and heterozygote plants derived from the *Ljcsld1-2* x wild type crosses. By contrast, a significantly decreased root hair length was observed in heterozygote plants of the *LjCSLD1*/*Ljcsld1-1* genotype (Fig. 7).

**Figure 7.**
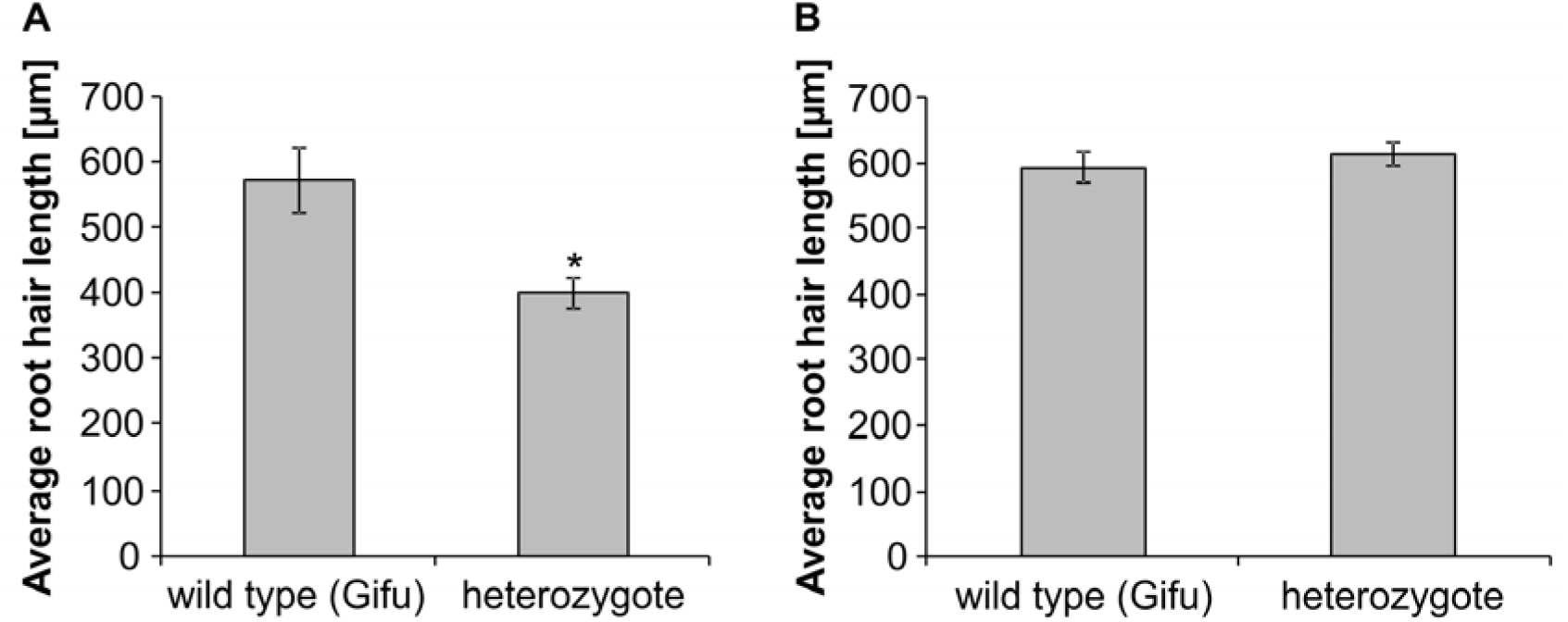
*Ljcsld1-1* allele exerts a semi-dominant effect on root hair growth. F2 progeny derived from crosses between *Ljcsld1-1* and Gifu (A) and *Ljcsld1-2* and Gifu (B) were genotyped and the root hair length was measured for homozygous wild-type (wild-type Gifu; n = 12 and 22 for A and B, respectively) and heterozygote individuals (n = 41 and 51). The values represent averages ± CI. Asterisk (*) denotes statistically significant difference from the wild-type, homozygous genotype, as determined using a Student’s *t*-test (P<0.05).

The observed differences between the short vs variable root hair phenotypes, including the thickness of cell walls, prompted the question of whether this correlates with the severity of symbiotic phenotypes. When analyzed 21 days after inoculation (dai) with *M. loti*, the *Ljcsdl1-1* line developed mostly uncolonized nodule primordia, while only a few small nodules colonized by rhizobia were formed, confirming our previous data (Karas et al., 2005). By contrast, all mutant lines with the variable root hair phenotype formed large, colonized nodules (Fig. 8).

**Figure 8.**
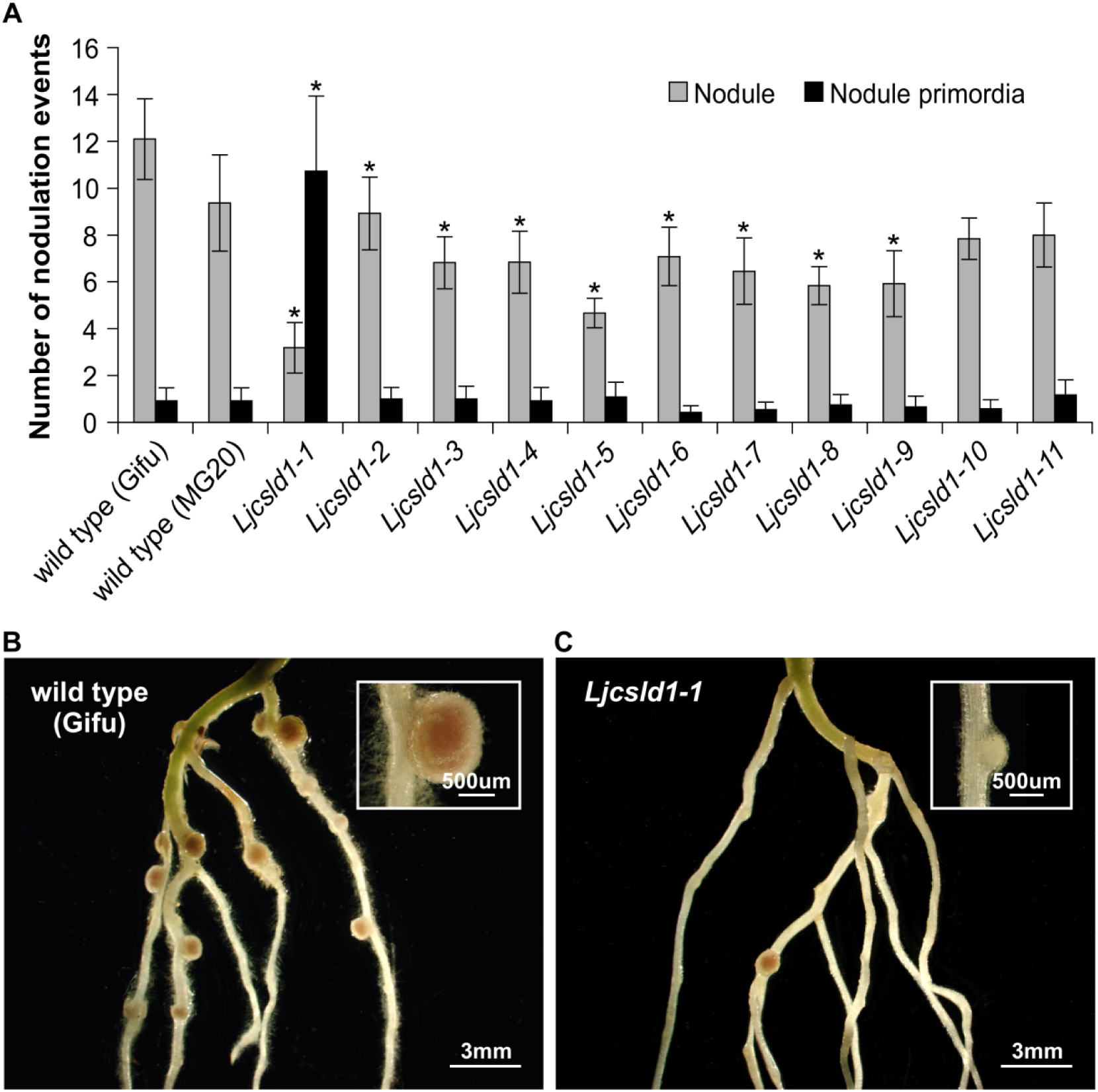
Symbiotic root phenotypes. Symbiotic root phenotypes of wild type *L. japonicus* Gifu and MG20 and the corresponding mutant plants carrying different *Ljcsld1* alleles; *Ljcsld1-1*, -*2*, -*4*, -*6* and -*7* alleles are in Gifu background while remaining alleles are in the MG20 ecotype. (A) Numbers of nodules and nodule primordia 21 days after inoculation (dai) with *M. loti* strain NZP2235; An average number ± 95% confidence interval is given for each genotype for n = 10 to 12. Asterisk (*) denotes statistically significant difference from the wild-type, homozygous genotype, as determined using a Student’s *t*-test (P<0.05). (B) Representative images of roots and root nodules for wild type (Gifu) and (C) *Ljcsld1-1* mutant. Note that all Gifu nodules are pink while the majority of the mutant nodules are underdeveloped and white in appearance (see inserts).

## Discussion

In this study, we identified a series of genetic lesions in the *L. japonicus Cellulose Synthase-Like D1* gene that cause root hair developmental defects. Two alleles, *Ljcsld1-1*, and *Ljcsld1-2*, complemented each other and five of the other nine alleles (*Ljcsld1-5, -7, -8, -9*, and *-10*) while the *Ljcsld1-3, -4, -6* and *-11* alleles failed to produce the wild-type root hair phenotype. Evidence indicates that CSLDs homodimerize or multimerize like their cellulose synthase counterparts (Taylor et al., 2003; Yin et al., 2011; Li et al., 2018). In this case, the intragenic complementation (i.e. complementation resulting from the formation of a functional multimeric protein from subunits produced by different mutant alleles of a gene) suggests that the mutant proteins encoded by *Ljcsld1-1* and *Ljcsld1-2* and those which they complement, retain partial functionality. This could be the result of dimerization/multimerization of the two mutant isoforms leading to a functional CSLD1 complex, or participation of both isoforms in a large heterocomplex with other CSLD subunits. The intragenic complementation and allele-specific phenotypes observed suggest modular functionality within the CSLD1 protein; the N-terminal domain can function independently of the C-terminal portion of the protein, and within the N-terminus, the Cys-rich region operates independently of the short proline-rich domain (PPTPD) which is mutated in *Ljcsld1-1*. The PPTPD region, which is specific to CSLDs, has been postulated to be involved in mediating protein-protein interactions (Delmer, 1999). Interestingly, the *Ljcsld1-2* mutation is in one of the seven of eight cysteine residues conserved with two zinc finger domains present in CESA1 (Supplemental Fig. S4B). For CESA, it was shown that the N-terminus contains conserved cysteine residues forming two zinc fingers and that oxidation of these cysteines promotes protein dimerization (Kurek et al., 2002). It is, therefore, possible that both of these mutations in the N-terminus affect homodimerization of CSLD1. In this case, the semi-dominant behavior of *Ljcsld1-1* could be explained through its improper interaction with wild-type subunits to form dimers with slightly reduced activity, while the unusually strong root hair phenotype of *Ljcsld1-1* homozygous mutants suggests not only that mutant dimers are inactive, but that they may interfere with any close homologs that may act redundantly. The *Ljcsld1-2* isoform, which acts recessively, may have reduced ability to form mutant homodimers/multimers but may interact with the wild type protein to form a functional complex. Interaction studies are needed to test these possibilities. Alternatively, the mutation in the *Ljcsld1-1* could be preventing appropriate degradation of the mutated CSLD1, resulting in overproduction of the corresponding cell wall component during the initial stage of growth of root hairs. This, in turn, would result in the thicker cell wall and possibly prevent the elongation of root hairs.

Another interesting observation was that intragenic complementation was only observed for specific alleles. The *Ljcsld1-1* and *Ljcsld1-2* N-terminal mutations were complemented by a subset of the mutations in the C-terminal catalytic domain or transmembrane domains. The inability of *Ljcsld1-3, -4, -6* and *-11* to complement *Ljcsld1-1* and *Ljcsld1-2* possibly reflects the severity of the mutations. For example, *Ljcsld1-6* and *- 11* both result in premature stop codons, and may, therefore, be null mutations. Notably, intragenic complementation has been observed for other CSLD and CESA proteins. The ZmCSLD1 proteins of *Zea mays* with G838D and W1184R substitutions in the catalytic and last transmembrane domain were shown to act in a complementary fashion (Li et al., 2018). Intragenic complementation was also reported for two alleles of *AtCesA3* (Pysh, 2015), but the mutations involved (G617E and G916E) were in a different region of the protein.

To find out if our discoveries could be applied to other species, we performed a cross-species complementation experiment which showed that AtCSLD2 and AtCSLD3 (KOJAK), required for root hair development in Arabidopsis (Bernal et al., 2008), are functionally equivalent to LjCSLD1. In *Populus trichocarpa* PtrCslD5 is considered a functional orthologue of AtCLSD3 and along with the highly homologous PtrCslD6, it is also involved in root hair development (Peng et al., 2019). Similarly, rice OsCSLD1 is also required for root hair morphogenesis (Kim et al., 2007).

Further, we analyzed cell wall composition using the entire roots of young wild-type and mutant seedlings. Using two different approaches, we showed a decrease of mannose across all mutant genotypes tested, which is intriguing as we observed significant symbiotic and non-symbiotic phenotypes between short and variable root hair mutants. It was previously suggested that mannose is enriched in soybean root hairs (Muszyński et al., 2015) therefore roots with short or missing root hairs would be predicted to have less of this component. On the other hand, glucose content in the cellulose fraction was increased in *Ljcsld1-1*, yet significantly diminished in all four variable root hair mutant lines. This increased value could well correlate with the root hair phenotype of *csld1* mutants which included thicker root hair cell walls, especially in *Ljcsld1-1* in which cell wall thickness had tripled, compared to wild type. An increased number of short cellulose fibrils could explain the higher glucose content. The increased thickness of root hair cell walls was also correlated with decreased nodulation of the mutants, again with *Ljcsld1-1* having the strongest phenotype. During rhizobial infection of root hairs, the plant remodels the cell wall at the point of infection by expressing a nodulation specific pectate-lyase, which is required for the formation of the infection thread (Xie et al., 2012). It is possible that the thicker cell wall of *Ljcsld1-1* presents a stronger barrier to infecting rhizobia. Alternatively, LjCSLD1 may be required for the formation of the infection thread cell wall. Another possibility is that non-growing root hairs are not susceptible to infection because they are unable to curl, which is required to capture bacterial and form a microcolony, from which IT will be initiated. By contrast, the mutant lines were found to be susceptible to arbuscular mycorrhizae (AM) fungi colonization (Novero and Bonfante, personal communication), consistent with the idea that root hairs are not the preferential penetration point for AM fungi.

Overall our allelic series provides a useful resource for investigating domains of the CSLD1 protein and their effects on cell wall formation, which can be scored by observing root hair development. Given that *L. japonicus* is amenable to hairy root transformation, it would be relatively simple to explore additional modifications in this gene. Knowledge acquired from this and future research using the newly discovered alleles should be able to be translated to other plant species and possibly to other cellulose synthase-like genes that function in different parts of the plant.

## Materials and Methods

### Plant material and growth conditions

*L. japonicus* root hair mutant lines S88-5A (*Ljsrh1)*, B12-IB (*Ljvrh1-1*), S49-AA (*Ljvrh1-2*), B69-A (*Ljvrh1-* and S36-1 were identified from a screen for genetic suppressors of the *L. japonicus* Gifu *har1-1* hypernodulation phenotype, as described (Karas et al., 2005; Murray et al., 2006b). Additionally, *L. japonicus* root hair mutants 210-226A, 212-010, 212-229, 01-0071, 212-164, and 212-447 were screened from ethyl methanesulfonate-treated M_2_ seeds of *L. japonicus* Miyakojima at RIKEN Plant Science Center and shared through NBRP Legume Base resource.

Unless otherwise stated, all plants were germinated and grown as described in (Karas et al., 2005). For cell wall analysis seeds (150 per replicate) were germinated and grown for 7 days on cellulose acetate film (Sterlitech Corporation, USA), moistened with water.

### Evaluation of root hair and symbiotic phenotypes

Symbiotic phenotypes were evaluated as described in (Karas et al., 2005). Length of root hairs of the F2 progeny derived from the crosses between wild type and either *Ljcsld1-1* or *Ljcsld1-2* were measured by photographing a 2 mm section of each root at the root hair mature zone (∼ 1 cm above the root tip). To make a measurement, two vertical axes were drawn, one at the base of the root and one at edge of the root hair tips such that the second axis included 95% or more of photographed root hairs. Each measurement is the average distance between the axes. For each root, measurements of root hairs were taken from each side of the root and averaged.

### Identification of full-length mRNA and coding regions

The *LjCSLD1* cDNA was amplified by RT-PCR of the total RNA derived from *L. japonicus* roots using the coding region-specific primers (CSLD1_cDNAF and CSLD1_cDNAR; see list of all primer sequences below). Rapid amplification of the 5’ and 3’ cDNA ends (RACE) was subsequently carried out by using the First Choice RLM-RACE kit from Ambion (USA), according to the manufacturer’s instructions. The full lenght *LjCSLD1* cDNA was reconstituted based on the obtained sequences. An additional set of two *LjCSLD1* mRNA-specific primers, positioned at the extreme 5’ and 3’ ends of the predicted full copy cDNA, were used (5’outer race, 5’inner race, 3’outer race, 3’inner race). Using total RNA from *L. japonicus* roots, RT-PCR was again performed and the resulting product was entirely sequenced.

### Expression analysis

Total RNA from *L. japonicus* tissues was isolated and converted into 1^st^ strand cDNA as described (Murray et al., 2007). To evaluate steady-state levels of the *LjCSLD1* mRNA in different *L. japonicus* tissues, the following PCR conditions were used: 5 min at 94°C, 30 cycles of 94°C for 30 sec, 60°C for 1 min, and 72°C for 30 sec, followed by 7 min at 72°C (primers: LjCSLD1expF, LjCSLD1expR); the *Ubiquitin* cDNA was amplified (primers: Ubi-F, Ubi-R; see primer list below) using similar PCR conditions.

### Transgenic hairy roots

The 8567 bp genomic fragment encompassing the *LjCSLD1* gene was amplified with primers LjCSLD1compF and LjCSLD1compR from BAC clone LjT30G11 (National Bio-Resource project website: https://www.legumebase.brc.miyazaki-u.ac.jp/index.jsp). The resulting cDNA was cloned into pDONR™221 vector (Invitrogen, USA) and subsequently moved to the pEarley303 destination vector. In addition, 6245, 4804, 4968, and 4675 bp genomic fragments encompassing LjCSLD1, AtCSLD2, AtCSLD3, and AtCSLA9, respectively, were cloned into the pEarley101 vector containing the *CaMV* 35S promoter. For histochemical analysis (promoter expression studies), a 2605 bp fragment of the *LjCSLD1* promoter was amplified using LjCSLD1prmt_F and LjCSLD1prmt_R, and cloned into the pBI101 binary vector. The resulting constructs were transferred into *Agrobacterium rhizogenes* AR10 strain and used to transform *L. japonicus* plants using the established protocol (Díaz et al., 2005). At least 20 independent plants were transformed for each construct analyzed. The resulting hairy roots were visually evaluated for the presence of root hairs. For the histochemical analysis of the reporter ß-glucuronidase (gusA) gene, transgenic hairy roots were stained as described in (Held et al., 2014).

### Cell wall analysis

At 7 days after germination, roots were harvested from a minimum of three biological replicates, cleared twice in a mixture of chloroform-methanol (1:1), and twice in acetone. Roots were subsequently air-dried, then weighed and analyzed in triplicate. Two independent methods (A and B) were used to analyze cell wall composition.

**A**)Roots were treated with 1M sulphuric acid (H_2_SO_4_) (105°C, 1 h) to hydrolyze non-cellulosic polymers. Hydrolysates were centrifuged to remove insoluble material, and after adding myo-inositol to each sample as an internal standard, the resulting soluble monosaccharides were derivatized to form alditol acetates (Blakeney et al., 1983) and analyzed by gas chromatography on an HP5880 gas chromatograph (Agilent, Santa Clara, CA) equipped with an SP-2330 column (0.25 mm x 30 m, Sigma Aldrich Canada, Oakville, ON) and a flame ionization detector. The insoluble residue was incubated in a mixture of acetic acid : nitric acid : water (8:2:1, 105°C, 1 h), then washed with water, and acetone and air dried (Updegraff, 1969). The pellet, was suspended in 67% H_2_SO_4_, solubilized by shaking (23°C, 1 h), diluted with water to a final concentration of 1M H_2_SO_4_ and assayed for cellulose content by the anthrone method as described by Updegraff (1969).

**B**)Cell walls from additional dried root samples were fractionated according to Fry (1988). Briefly, root tissue was extracted with 70% (v/v) ethanol (70°C, 1 h), and the alcohol insoluble residue was sequentially treated with DMSO : water (9:1, 23 °C, 16 h) to remove starch, 0.5% (w/v) ammonium oxalate (105°C, 1 h followed by 4°C, 16 h), to extract pectins and 4M KOH containing 0.1% (w/v) NaBH_4_ (23°C, 2 x 1 h), to isolate hemicelluloses.

Uronic acids (pectin) in the ammonium oxalate soluble supernatant were assayed based on the previously described method (Blumenkrantz and Asboe-Hansen, 1973). To 1 volume of supernatant, 5 volumes of 0.5% (w/v) borax solution in concentrated H_2_SO_4_ was added, mixed well, incubated (105°C, 5 min), then cooled in a water bath at 23°C. Absorbance at 520 nm (absorbance A) was measured. Subsequently, 1/10 volume, based on the original supernatant volume, of a 0.15% (w/v) solution of 3-phenylphenol in 1N NaOH was added, mixed well and incubated (23°C, 5 min) before re-reading the absorbance at 520 nm (absorbance B). Uronic acid content was determined by subtracting absorbance A from absorbance B and comparing to a standard curve prepared using galacturonic acid.

Hemicelluloses in the KOH supernatant were assayed by neutralizing the extract with acetic acid, dialyzing against water to remove salts and hydrolyzing to component monosaccharides in 1M H_2_SO_4_ (105°C, 1 h). Monosaccharides were converted to alditol acetates and quantified by the method of (Blakeney et al., 1983) as described in method **A**, above.

### Evaluation of root hair cell wall thickness

Roots segments were excised, fixed in glutaraldehyde and osmium tetroxide and infiltrated in Epon/Araldite resin. In order to be able to obtain complete (from the tip to the insertion point) longitudinal sections of root hairs, we used a “flat embedding” procedure.

A glass microscope slide was covered by a thick teflon layer. Root segments were subsequently placed on the slide with a small drop of resin and covered with a Teflon coverslip. The slides were then incubated (60°C, 24 h) to allow resin polymerization. After removal of the Teflon coverslips, the “slides” were examined under a light microscope and the best points were selected and cut in order to obtain semi-thin and thin sections.

To verify the different cell wall thickness in WT, variable and short root hair mutants, we selected five short root hairs, six WT root hairs, seven variable “Short Type” root hairs and ten variable “WT Type” root hairs (only longitudinal sections were considered). For each root hair, we obtained TEM picture at 39000X magnification from the same region (corresponding to the middle part of the root hair, avoiding the apex and the basal zone) and we measured the cell wall thickness at four different points for each TEM pictures. For each TEM picture, we calculated the mean value of the four measurements. The detection of the fibrillar organization was performed by treating the thin sections with the periodic acid-thiocarbohydrazide-silver proteinate (PATAg) reagent for polysaccharide localization (Roland and Vian, 1991)

## List of primers used in this study

All sequences are presented in a 5’ to 3’ orientation.

*Map-based cloning*

JM003_F TCGGAGACAGAAGGCATCTT

JM003_R GCTTGTCTGGGAAGCTGTTC

JM010_F ACTATACGTCGCACCAAACG

JM010_R GCCATGACTGGAGCAGAAAC

*Amplification of LjCSLD1 cDNA*

CSLD1_cDNAF TGACAGTGAGCTGGGAAGTG

CSLD1_cDNAR AGCACCCAAAAGCTGAAGAA

*RACE*

3’outer race GTCGATGACGAGTTTGCTGA

3’inner race CAGCACCATACCTCAGTGGA

5’outer race GCTGTTGAATCCTCCGGTAA

5’inner race TTGGTTATCAGGGGTTGGTG

*Evaluation of LjCSLD1 in L. japonicus tissues*

LjCSLD1expF CCGTTTGCTAGGTGGTGTTT

LjCSLD1expR TCACCATCAAACGTTCCTGA

Ubi-F TTCACCTTGTGCTCCGTCTTC

Ubi-R AACAACAGAACACACAGACAATCC

*Construction of LjCSLD1p::GUS plasmid*

LjCSLD1prmt_F TTTTTGTTTTGGTCAACAGCAG

LjCSLD1prmt_R TACACTGGCACACGGAGAGA

*Construction of LjCSLD1p::LjCSLS1 plasmid*

LjCSLD1compF TTGGTCCACAAACAGCTGAA

LjCSLD1compR TCACCATCAAACGTTCCTGA

*Construction of 35Sp::LjCSLS1, AtCSLA9, AtCSLD2, AtCSLD3 plasmids*

LJCSLD1_35S_F TCCGTGTGCCAGTGTATGTT

LJCSLD1_35S_R GAGACAACAAAAAGCCTTGGA

ATCSLA9_F TCCTTTTTCCCGACAATCTG

ATCSLA9_R TGCCTTCAAGGAATCTGAAAA

ATCSLD2_F AAAGATGTGTGGGCTTTTCG

ATCSLD2_R TCCACCCAATCTTGTTCCAT

ATCSLD3_F TGTCCGAAAACCTCAACACA

ATCSLD3_R CCCGTTTTGTTTCCTTTTCC

## List of author contributions

B.J.K and K.S. conceived the original screening and research plans; B.J.K and K.S. supervised the experiments; B.J.K, L.R. and L.A performed most of the experiments; B.J.K, L.R., M.N., L.A., J.D.M, P.B., K.S. designed the experiments and analyzed the data; B.J.K and K.S. conceived the project and wrote the article with contributions from all of the authors; B.J.K. and K.S., supervised and complemented the writing. S.I, M.N., and T.S., provided seeds for the following mutant lines: 210-226A, 212-010, 212-229, 01-0071, 212-164, and 212-447; S.S. provided L. japonicus genome sequence data; P.B. and M.N performed the transmission electron-microscopy experiment; B.J.K agrees to serve as the author responsible for contact and ensures communication.

## Acknowledgments

We thank Alex Molnar for the preparation of the figures, Dr. Jenny Mortimer and Paul Dupree for additional work on cell wall composition that is not included in this manuscript, Dr. Jenny Mortimer for providing helpful comments on the manuscript.

